# Brain fingerprinting using EEG graph inference

**DOI:** 10.1101/2023.03.11.532201

**Authors:** Maliheh Miri, Vahid Abootalebi, Enrico Amico, Hamid Saeedi-Sourck, Dimitri Van De Ville, Hamid Behjat

## Abstract

Taking advantage of the human brain functional connectome as an individual’s fingerprint has attracted great research in recent years. Conventionally, Pearson correlation between regional time-courses is used as a pairwise measure for each edge weight of the connectome. Building upon recent advances in graph signal processing, we propose here to estimate the graph structure as a whole by considering all time-courses at once. Using data from two publicly available datasets, we show the superior performance of such learned brain graphs over correlation-based functional connectomes in characterizing an individual.

## 1. Introduction

A growing number of studies have demonstrated that functional connectomes are subject-specific and can thus be considered as brain fingerprints; i.e., capable to identify an individual within a population, in health [1] and disease [2], [3]. The conventional approach is to construct the functional connectome (FC) by considering brain regions as vertices and a pairwise measure of statistical dependence (i.e., Pearson’s correlation coefficient) between the regional time-courses of pairs of regions as edge weight. The fingerprinting potential of FC has been investigated using different neuroimaging modalities, namely electroencephalography (EEG) [4], [5], magnetoencephalography (MEG) [6], [7], and functional magnetic resonance imaging (fMRI) [1], [8]. All these studies have helped in advancing towards single-subject level inferences from brain connectivity data, i.e., by capitalizing on individual properties of functional network organization across different cognitive tasks and resting state [9], [10], or by relating individual connectome features to behavioral and demographic scores [1], [6], [7], [9]. However, the conventional FC captures not only the statistical dependence between neural activity but also that of underlying noise sources. Furthermore, functional connectomes by construction only provide a pairwise representation of the brain dynamics, e.g., by looking at the brain as a composition of dyads. Beneficial because of its simplicity, this assumption limits the investigation of the individual features arising from human brain networks. Therefore, FC denoising remedies have been proposed based on principal component-based reconstruction [9] or eigenspace embedding [10], each of which require the learning of latent space-based FCs from a population of subjects. Here we propose an alternative strategy, that is, to directly infer a sparse graph structure [11] from an individual’s EEG data—thus, an alternative to a functional connectome—using principles from graph signal processing (GSP) [12], [13]. The fundamental objective of the learning strategy is to infer a graph on which brain maps are seen as smooth signals—that is, the majority of signal energy lies at the lower end of the graph spectrum, which is an intrinsic property of functional brain organization supported by prior results in brain GSP [14]–[19]. We treat the adjacency matrix of learned graphs as an alternative to FC matrices for fingerprinting. A closely related method has already shown promising results on fMRI data [8]. Here we focus on EEG data, leveraging our recently proposed method for inferring brain graphs from EEG data [20]. This method results in a graph that captures subtle spatial relations between EEG electrodes, in particular, their instantaneous spatial profiles, such that EEG maps can be seen as smooth functions on the resulting graphs. Using two open-access EEG datasets, we show that learned graphs provide superior fingerprinting performance over standard functional connectomes. The remainder of the paper is organized as follows. Section II gives an overview of fundamental concepts and the proposed method. Section III presents the experimental setup, the results, and a discussion. Our concluding remarks are presented in Section IV.

## II. Methods

### A. Graph Learning via Enforcing Graph Signal Smoothness

Let *G* denote an undirected, weighted graph with no self-loops, represented by an *N × N* adjacency matrix **A**, where *N* denotes the number of vertices, with elements *A*_*i,j*_ representing a measure of similarity of connection strength between vertices *i* and *j*, with *A*_*i,i*_ = 0. A graph signal **f** *∈* ℝ^*N*^ can be seen as a function residing on the vertices of the graph, whose *i*-th component is the signal value at the *i*-th vertex. The graph degree matrix **D** is given as *D*_*i,i*_ = *A*_*i,j*_, and the graph’s combinatorial Laplacian matrix **L** is given as **L** = **D** *−* **A**. Eigendecomposing **L** gives **L** = **UΛU**^*T*^ where **U** = [**u**_1_, **u**_2_, …, **u**_*N*_] concatenates the orthonormal eigenvectors in its columns, and **Λ** is the diagonal matrix of the corresponding eigenvalues **Λ** = *diag*(*λ*_1_, …, *λ*_*N*_). The eigenvalues define the graph Laplacian spectrum. Each eigenvalue represents the extent of variability of the associated eigenvector relative to the graph structure. In particular, the variability of a graph signal **f** —where graph Laplacian eigenvectors may also be seen as graph signals—can be quantified using a measure of total variation (TV) as [21]:

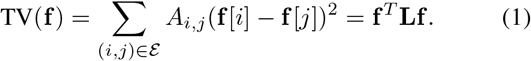

Given two graph signals **f**_1_ and **f**_2_, if TV(**f**_1_) *<* TV(**f**_2_), then **f**_1_ is a smoother graph signal than **f**_2_ as it exhibits less variability on the graph.

A major paradigm in graph structure learning exploits this notion of smoothness to infer a sparse graph from a given set of observations [11], [22]. Specifically, a graph combinatorial Laplacian matrix can be inferred via the following optimization [22]:

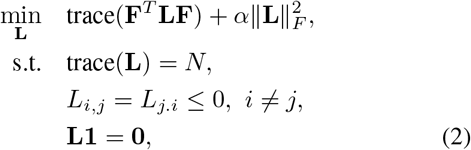

where **F** is an *N × M* matrix of *M* graph signals, *α* is regularization parameter, *∥·∥*_*F*_ denotes the Frobenius norm and **1** = [1, …, 1]^*T*^ . Minimizing the first term guarantees smoothness of the signals on the learned graph, which can be seen via invoking (1): trace(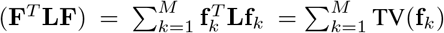

The Frobenius norm controls graph sparsity by shrinking edge weights. The imposed constraints in the optimization ensure finding a valid Laplacian matrix. Noting that 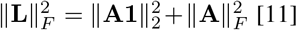 [11], replacing the *ℓ*_2_-norm with a logarithmic barrier, the optimization in (2) can be solved more efficiently via a more general-purpose formulation with respect to the graph’s adjacency matrix [11]:

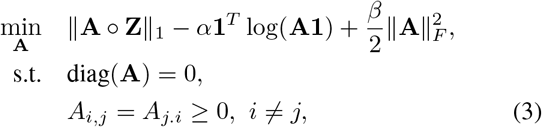

where **Z** is an *N × N* matrix with elements *Z*_*i,j*_ = *∥***F**_*i*,:_ *−* **F**_*j*,:_*∥*_2_, i.e., Euclidean distance between signal values on electrodes *i* and *j*. The first term in (3) enforces the smoothness constraint in similar way as in the first term in (2), which is based on the equivalence trace(**F**^*T*^ **LF**) = 0.5*∥***A** *◦* **Z***∥*_1_, where *◦* is the Hadamard product [20]. Intuitively, if smooth graph signals reside on well-connected vertices, it is expected that these vertices have smaller distances *Z*_*i,j*_. The second term enforces graph degrees to be positive and improves the overall connectivity of the graph. *α* and *β* are regularization parameters, the constraints guarantee to obtain a valid adjacency matrix, and the last term controls the graph sparsity.

### B. Differential identifiability

The underlying assumption in the maximization of human functional connectome fingerprinting is that the connectivity profiles related to the same subject should be more similar than they are across different subjects. In order to evaluate our method, we utilized the approach provided in [9], in which, the identifiability capability is quantified by defining the concept of “level of identifiability” on a set of FCs. In particular, we consider a generalized definition of identifiability for any desired feature set, not just FC. Given *S* subjects, let 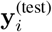 and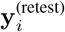 denote the test and the retest feature vectors of subject *i* where *i* = 1, …, *S* and *S* denotes the number of subjects. An *S × S* identifiability matrix **M** for the given feature set can then be defined, with elements *M*_*k,l*_ equal to the Pearson correlation between 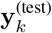 and 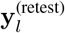 Note that this matrix is not symmetric, because of the test/retest encoding. Given **M**, a scalar value *I*_self_ can be defined as the average of the main diagonal elements—the correlation between test and retest feature sets of same subjects—that indicates the average self-similarity across subjects. Accordingly, a scalar value *I*_others_ can be defined as the average of the off-diagonal elements—the correlation between different subjects’ test and retest feature sets—that quantifies the average cross-subject similarity based on the feature set at hand. A third scalar value is defined as *I*_diff_ = (*I*_self_ -*I*_others_)*×*100, providing a robust group level-estimate of identifiability at the individual connectome level, where a higher value indicates greater individual fingerprinting [9].

It is insightful to quantify the influence of individual graph edges to fingerprinting. Considering that subject identity is determined based on the largest value of the Pearson’s correlation between the test and all the retest feature sets, the contribution of each edge in computing the correlation can be treated as edge significance [1], [9]. In particular, let 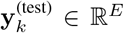 and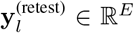 denote two z-score normalized feature vectors, where *k, l* = 1, …, *S*, and *E* = (*N* ^2^ *− N*)*/*2. A measure of fingerprinting strength of each edge can be defined as:

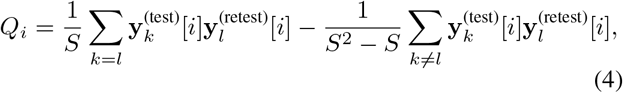

where *i* = 1, …, *E* denotes edge index. The higher the value of *Q*_*i*_, the higher the impact of edge *i* overall along the population in fingerprinting. Consequently, using *{Q*_*i*_*}*_*i*=1,…,*E*_ a measure of nodal fingerprinting strength can be obtained for each graph node *n* = 1, …, *N* by averaging *Q*_*i*_ values associated to each node.

### C. Datasets

We used EEG data from two publicly available datasets: i) motor-imagery task data from the BCI Competition III-Dataset IVa^1^ [23], and ii) a resting-state data of 109 subejcts ^2^ [24], [25]. In the remainder of this paper we denote these two datasets as Dataset-1 and Dataset-2, respectively. Dataset1 contains EEG signals from five subjects, acquired using 118 electrodes at a sampling rate of 100 Hz. The data for each subject consists of 280 trials, for two different classes of motor imagery tasks (right hand and right foot), 140 trials per class; each trial lasted for 3.5 seconds, each of which contained rest intervals of varying lengths from 1.75 to 2.25 seconds at the end. Dataset-2 contains EEG signals from 109 subjects, by 64 electrodes using BCI2000 system [24] with a sampling rate of 160 Hz. Each subject performed fourteen different experimental runs, including two one-minute restingstate runs (in “eyes open” and “eyes closed” modes), and three two-minute runs of real and imagery movements. In order to prevent the possible bias of different sessions and tasks, we used the resting-state recording in both eyes-open and eyesclosed conditions for our fingerprinting purposes.

### D. EEG-based fingerprinting

For each subject in each of the two datasets, we infer a Functional Learned Graph (FLG), using different sets of EEG maps from each subject; detail on sets given below. In each graph, vertices represent EEG electrodes (same definition across subjects for each dataset), whereas edges and their weights were derived by estimating the graph’s weighted adjacency matrix using the optimisation in (3). As a means of comparison, using the EEG data used for each graph learning setting, we also defined a fully connected FC Graph (FCG) in which edge weights were defined based on the absolute value of the Pearson’s correlation between pairs of electrodes. In Dataset-1, we extract from each trial the 0.5-2.5 second interval after the visual cue [26], and treat these as a task-active trial, and we treat the 1.75 seconds interval after the trials as a rest trial. Given that motor activity (real or imagined) modulates the sensorimotor mu rhythm (8-13 Hz) and beta rhythm (13-30 Hz), we bandpass filtered the extracted signals with a third-order Butterworth filter to retain 8-30 Hz frequencies. We compared four different settings for the set of EEG signals used for building FCG and FLG, namely, using trials of both tasks, just task 1, just task 2, and rest trials. In each setting, we treat half of the signals as the test set and the other half as the retest set. In Dataset-2, the resting-state EEG signals from two modes were temporally bandpass filtered into six canonical frequency bands: delta (0.5-4 Hz), theta (4-8 Hz), alpha (8-13 Hz), low beta (13-20 Hz), high beta (20-30 Hz) and gamma (30-50 Hz). In each band, the first 30 seconds were treated as test set, whereas the remaining 30 seconds were treated as retest set. FLG and FCG were derived for the test and retest sets of each subject and the vectorized version of the upper triangular part of them were considered as feature sets.

## III. Results

### EEG functional learned graph

Fig. 1 shows representative graphs and their related adjacency matrices obtained using FLG and FCG. The FLG is a much sparser representation compared to the FCG, which is a complete graph. This property can be better observed by inspecting the corresponding adjacency matrices that are shown in Fig. 1(c); given that the matrices are symmetric, only the lower/upper triangular parts of them are shown.

**Fig. 1:**
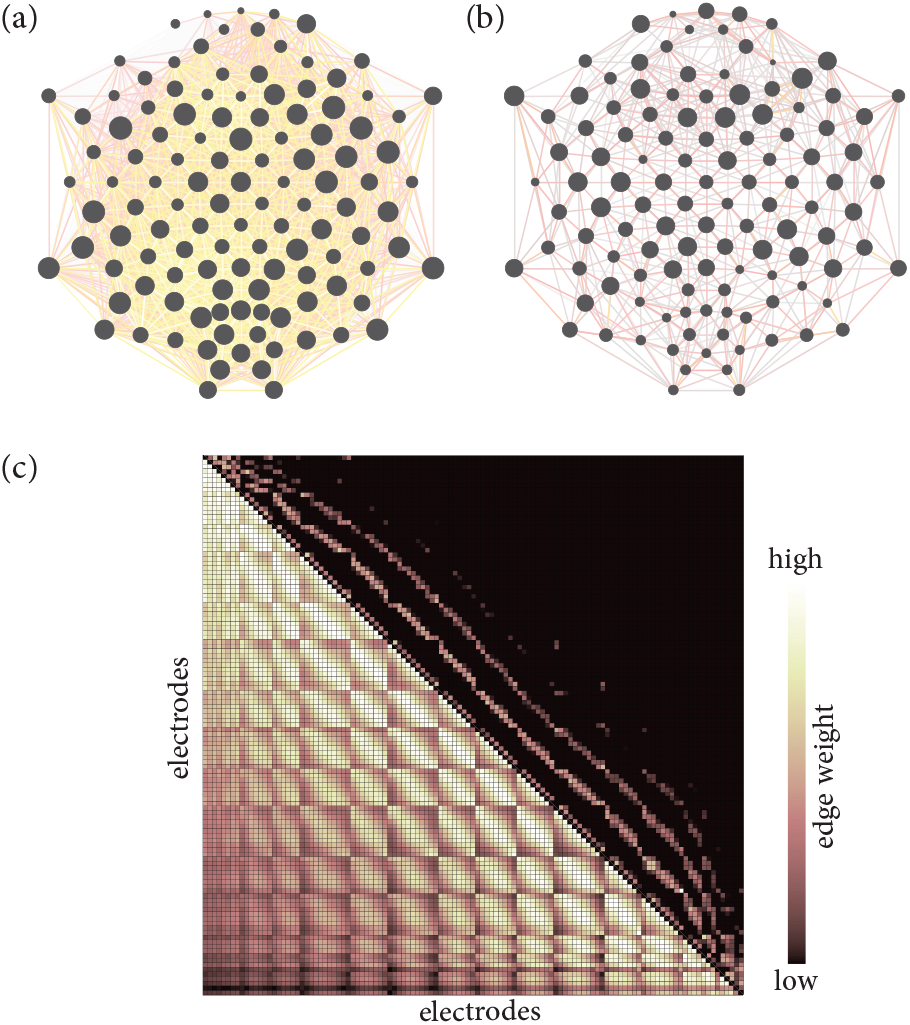
A representative subject-specific (a) FCG, and (b) FLG. (c) Adjacency matrices of the graphs shown in (a) and (b); the lower triangular section is for the FCG whereas the upper triangular section is for the FLG.

**Fig. 2:**
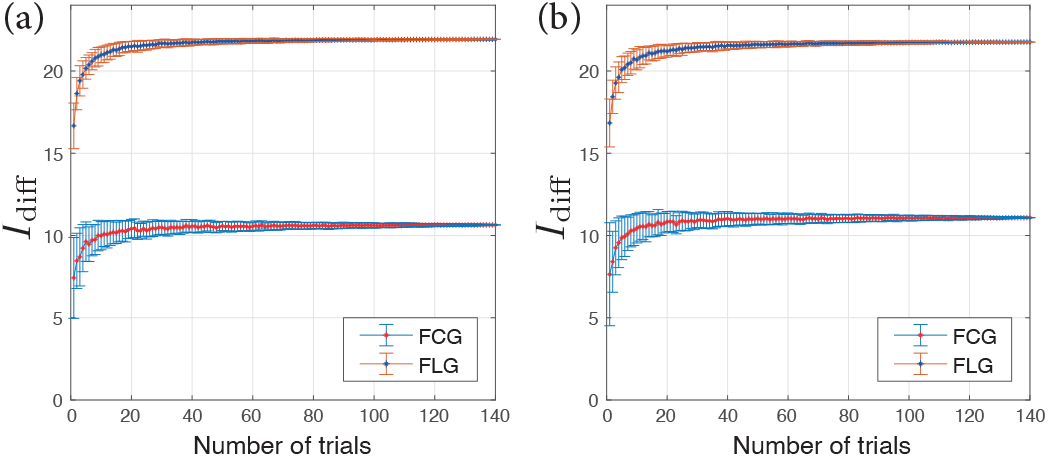
Differential identifiability power of FCG and FLG as a function of the number of trials used to build and learn the graphs, respectively; results on (a) both tasks, and (b) rest trials of Database-1. The error bars show the standard deviation across 100 bootstrap iterations.

### Fingerprinting power of FLG over FCG

TABLE I and TABLE II show fingerprinting results using FCG and FLG on Dataset-1 and Dataset-2, respectively. The results indicate the superior performance of learned graphs compared to conventional correlation-based connectivity graphs in both *I*_diff_ and accuracy measures. In Dataset-1, for all four settings, *I*_diff_ values obtained from using FLG are substantially higher than the values of FCG, and subjects are identified with absolute accuracy using both FLG and FCG. Similarly, in Dataset-2, for all frequency bands, FLG outperforms FCG. Specifically, all subjects are identified in the Eyes Open mode in the last four bands, as well as in the alpha and low beta bands in the Eyes Closed mode, which outperform the results of the state-of-the-art methods [5], [27].

**TABLE I:**
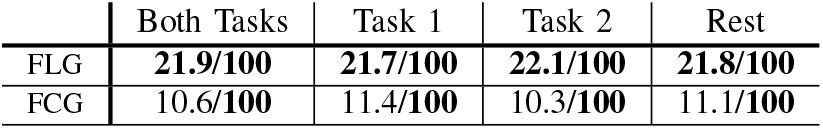
Fingerprinting *I*_diff_ (%)/accuracy(%) on Dataset-1.

|  | Both Tasks | Task 1 | Task 2 | Rest |
| --- | --- | --- | --- | --- |
| FLG | **21.9/100** | **21.7/100** | **22.1/100** | **21.8/100** |
| FCG | 10.6/100 | 11.4/100 | 10.3/100 | 11.1/100 |

**TABLE II:**
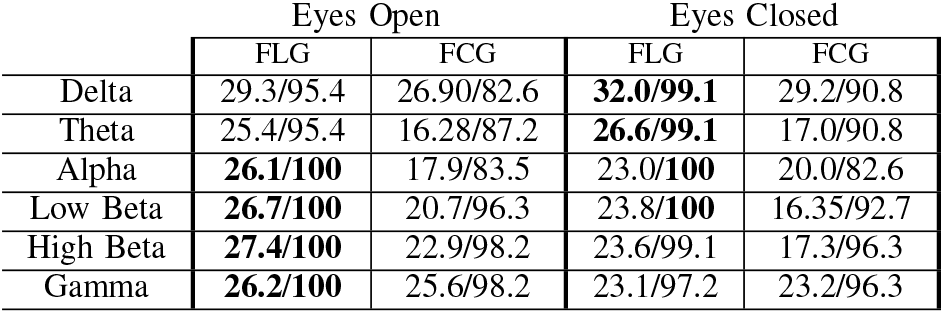
Fingerprinting *I*_diff_ (%)/accuracy(%) on Dataset-2.

|  | Eyes Open |  | Eyes Closed |  |
| --- | --- | --- | --- | --- |
|  | FLG | FCG | FLG | FCG |
| Delta | 29.3/95.4 | 26.90/82.6 | **32.0/99.1** | 29.2/90.8 |
| Theta | 25.4/95.4 | 16.28/87.2 | **26.6/99.1** | 17.0/90.8 |
| Alpha | **26.1/100** | 17.9/83.5 | 23.0/100 | 20.0/82.6 |
| Low Beta | **26.7/100** | 20.7/96.3 | 23.8/100 | 16.35/92.7 |
| High Beta | **27.4/100** | 22.9/98.2 | 23.6/99.1 | 17.3/96.3 |
| Gamma | **26.2/100** | 25.6/98.2 | 23.1/97.2 | 23.2/96.3 |

### The impact of EEG duration on fingerprinting

In order to investigate the effect of the size of EEG time series on fingerprinting, we performed identification using different numbers of trials in Dataset-1 and different lengths of epochs in Dataset-2. In particular, in Dataset-1, we used 280 trials from both tasks and rest (half of them for test and half for retest sets), and differential identifiability was obtained as the average of 100 bootstraps for each instance of number of trials; that is, in each bootstrap iteration we randomly selected *n* trials and then derived the FCG and the FLG. In Dataset-2, given that there were no trials available, we varied the length of the EEG epochs used for deriving the graphs from 2 to 30 seconds for both the test and retest sets. The results are presented in Fig. 3. On both Eyes Open and Eyes Closed rest data, fingerprinting performance enhances as a function of length of epochs in all frequency bands with the exception of the Gamma band for which the performance reaches a light plateau after around eight seconds. FLG clearly outperform FCG across all setting, once again, with the exception of the Gamma bound for which the performances are alsmot identical; the largest difference in performance between FLG and FCG is within the Theta band, a pattern that is consistent along different epoch lengths.

**Fig. 3:**
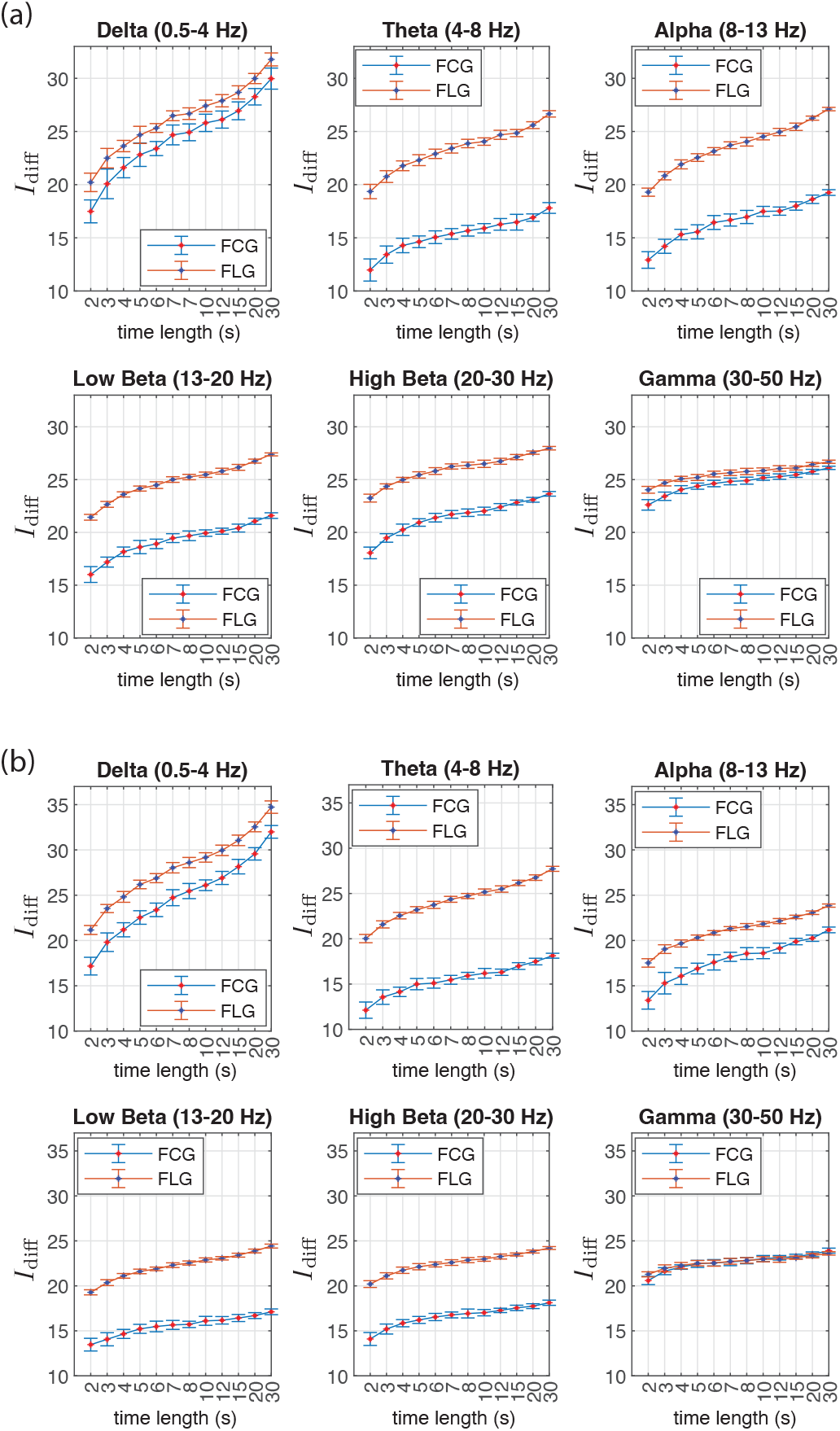
Differential identifiability power of FCG and FLG as a function of the length of epochs used to derive the graphs for six frequency bands on the (a) Eyes Open and (b) Eyes Closed datasets; results on Database-2. The error bars show the standard deviation across 30 bootstrap iterations.

### Significance of individual graph edges in fingerprinting

Fig. 4 shows electrode maps specifying the significance of individual electrodes/cortical-regions for fingerprinting, where significance was computed using the method described in Section II-B. In all six frequency bands, results show spatially localized patterns that are mainly in the frontal lobe, areas of the fronto-parietal network that have previously also been linked to fingerprinting in related work on fMRI data [1], [28].

**Fig. 4:**
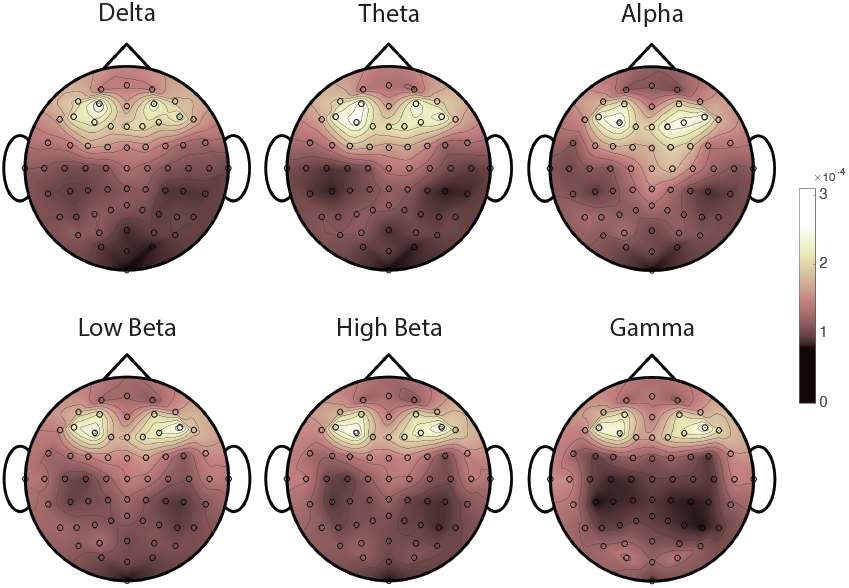
Significance of brain regions for fingerprinting.

## Conclusions

We proposed the use of brain graphs learned from EEG data for the purpose of identifying individuals, as an alternative to conventional correlation-based functional connectomes. Results showed superior performance of the proposed graphs over FC graphs on two datasets. The fingerprinting potential of learned EEG graphs may be further explored in future work with EEG source-reconstructed functional connectomes, as well as to leverage latent space embedding schemes [9], [10]. Future work should also explore the potentials of this methodology in other neuroimaging modalities (e.g. MEG [6], fMRI [9]), or to study the effect of brain stimulation on individual identifiability (e.g., after Transcranial Magnetic Stimulation). Finally, the eigenbases of EEG learned graphs may prove beneficial in providing a more compact feature space [29] for fingerprinting, in particular, their distance to a common harmonic basis [30], [31] derived from a large sample of subjects.

## Footnotes

1 https://www.bbci.de

2 https://physionet.org/content/eegmmidb/1.0.0/

## References

[1] E. S. Finn, X. Shen, D. Scheinost, M. D. Rosenberg, J. Huang, M. M. Chun, X. Papademetris, and R. T. Constable, “Functional connectome fingerprinting: identifying individuals using patterns of brain connectivity,” Nature neuroscience, vol. 18, no. 11, pp. 1664–1671, 2015.

[2] P. Sorrentino, R. Rucco, A. Lardone, M. Liparoti, E. T. Lopez, C. Cavaliere, A. Soricelli, V. Jirsa, G. Sorrentino, and E. Amico, “Clinical connectome fingerprints of cognitive decline,” NeuroImage, vol. 238, p. 118253, 2021.

[3] S. Stampacchia, S. Asadi, S. Tomczyk, F. Ribaldi, M. Scheffler, K.-O. Lövblad, M. Pievani, G. B. Frisoni, V. Garibotto, and E. Amico, “Fingerprinting of brain disease: Connectome identifiability in cognitive decline and neurodegeneration,” bioRxiv, pp. 2022–02, 2022.

[4] M. Fraschini, S. M. Pani, L. Didaci, and G. L. Marcialis, “Robustness of functional connectivity metrics for EEG-based personal identification over task-induced intra-class and inter-class variations,” Pattern Recognition Letters, vol. 125, pp. 49–54, 2019.

[5] M. Demuru and M. Fraschini, “EEG fingerprinting: Subject-specific signature based on the aperiodic component of power spectrum,” Computers in Biology and Medicine, vol. 120, p. 103748, 2020.

[6] E. Sareen, S. Zahar, D. Van De Ville, A. Gupta, A. Griffa, and E. Amico, “Exploring MEG brain fingerprints: evaluation, pitfalls, and interpretations,” Neuroimage, vol. 240, p. 118331, 2021.

[7] J. da Silva Castanheira, H. D. Orozco Perez, B. Misic, and S. Baillet, “Brief segments of neurophysiological activity enable individual differentiation,” Nature communications, vol. 12, no. 1, p. 5713, 2021.

[8] S. Gao, X. Xia, D. Scheinost, and G. Mishne, “Smooth graph learning for functional connectivity estimation,” Neuroimage, vol. 239, p. 118289, 2021.

[9] E. Amico and J. Goñi, “The quest for identifiability in human functional connectomes,” Scientific Reports, vol. 8, no. 1, pp. 1–14, 2018.

[10] K. Abbas, E. Amico, D. O. Svaldi, U. Tipnis, D. A. Duong-Tran, M. Liu, M. Rajapandian, J. Harezlak, B. M. Ances, and J. Goñi, “GEFF: Graph embedding for functional fingerprinting,” NeuroImage, vol. 221, p. 117181, 2020.

[11] V. Kalofolias, “How to learn a graph from smooth signals,” in Artificial Intelligence and Statistics. PMLR, 2016, pp. 920–929.

[12] A. Ortega, P. Frossard, J. Kovačević, J. M. F. Moura, and P. Vandergheynst, “Graph signal processing: Overview, challenges, and applications,” Proc. IEEE, vol. 106, no. 5, pp. 808–828, 2018.

[13] W. Huang, T. A. W. Bolton, J. D. Medaglia, D. S. Bassett, A. Ribeiro, and D. V. D. Ville, “A graph signal processing perspective on functional brain imaging,” Proceedings of the IEEE, vol. 106, no. 5, pp. 868–885, 2018.

[14] M. Miri, V. Abootalebi, and H. Behjat, “Enhanced motor imagery-based EEG classification using a discriminative graph fourier subspace,” in 2022 IEEE 19th International Symposium on Biomedical Imaging (ISBI). IEEE, 2022, pp. 1–5.

[15] K. Glomb, J. R. Queralt, D. Pascucci, M. Defferrard, S. Tourbier, M. Carboni, M. Rubega, S. Vulliemoz, G. Plomp, and P. Hagmann, “Connectome spectral analysis to track EEG task dynamics on a subsecond scale,” NeuroImage, vol. 221, p. 117137, 2020.

[16] H. Behjat, A. Tarun, D. Abramian, M. Larsson, and D. Van De Ville, “Voxel-wise brain graphs from diffusion mri: Intrinsic eigenspace dimensionality and application to fmri,” bioRxiv, pp. 2022–09, 2022.

[17] H. Behjat and M. Larsson, “Spectral characterization of functional MRI data on voxel-resolution cortical graphs,” in Proc. IEEE Int. Symp. Biomed. Imaging. IEEE, 2020, pp. 558–562.

[18] H. Behjat, N. Leonardi, L. Sörnmo, and D. Van De Ville, “Anatomically-adapted graph wavelets for improved group-level fMRI activation mapping,” Neuroimage, vol. 123, pp. 185–199, 2015.

[19] S. Atasoy, L. Roseman, M. Kaelen, M. L. Kringelbach, G. Deco, and R. L. Carhart-Harris, “Connectome-harmonic decomposition of human brain activity reveals dynamical repertoire re-organization under LSD,” Scientific reports, vol. 7, no. 1, pp. 1–18, 2017.

[20] M. Miri, V. Abootalebi, H. Saeedi-Sourck, D. Van De Ville, and H. Behjat, “Spectral Representation of EEG Data using Learned Graphs with Application to Motor Imagery Decoding,” bioRxiv, pp. 2022–08, 2022.

[21] U. Von Luxburg, “A tutorial on spectral clustering,” Stat. Comput., vol. 17, pp. 395–416, 2007.

[22] X. Dong, D. Thanou, P. Frossard, and P. Vandergheynst, “Learning Laplacian matrix in smooth graph signal representations,” IEEE Trans. Signal Process., vol. 64, no. 23, pp. 6160–6173, 2016.

[23] B. Blankertz, K.-R. Muller, D. J. Krusienski, G. Schalk, J. R. Wolpaw, A. Schlogl, G. Pfurtscheller, J. R. Millan, M. Schroder, and N. Birbaumer, “The BCI competition III: Validating alternative approaches to actual BCI problems,” IEEE Trans. Neural Syst. Rehabil. Eng., vol. 14, no. 2, pp. 153–159, 2006.

[24] G. Schalk, D. J. McFarland, T. Hinterberger, N. Birbaumer, and J. R. Wolpaw, “BCI2000: a general-purpose brain-computer interface (BCI) system,” IEEE Transactions on biomedical engineering, vol. 51, no. 6, pp. 1034–1043, 2004.

[25] A. L. Goldberger, L. A. Amaral, L. Glass, J. M. Hausdorff, P. C. Ivanov, R. G. Mark, J. E. Mietus, G. B. Moody, C.-K. Peng, and H. E. Stanley, “PhysioBank, PhysioToolkit, and PhysioNet: components of a new research resource for complex physiologic signals,” circulation, vol. 101, no. 23, pp. e215–e220, 2000.

[26] K. Georgiadis, D. A. Adamos, S. Nikolopoulos, N. Laskaris, and I. Kompatsiaris, “Covariation informed graph Slepians for motor imagery decoding,” IEEE Trans. Neural Syst. Rehabil. Eng., vol. 29, pp. 340–349, 2021.

[27] J. C. Monsy and A. P. Vinod, “EEG-based biometric identification using frequency-weighted power feature,” IET Biometrics, vol. 9, no. 6, pp. 251–258, 2020.

[28] A. Griffa, E. Amico, R. Liégeois, D. Van De Ville, and M. G. Preti, “Brain structure-function coupling provides signatures for task decoding and individual fingerprinting,” Neuroimage, vol. 250, p. 118970, 2022.

[29] S. Maghsadhagh, J. L. D. da Rocha, J. Benner, P. Schneider, N. Golestani, and H. Behjat, “A discriminative characterization of Heschl’s gyrus morphology using spectral graph features,” in Proc. IEEE Int. Conf. Eng. Med. Biol. Soc. IEEE, 2021, pp. 3577–3581.

[30] J. Chen, G. Han, H. Cai, D. Yang, P. J. Laurienti, M. Styner, and G. Wu, “Learning common harmonic waves on Stiefel manifold–a new mathematical approach for brain network analyses,” IEEE transactions on medical imaging, vol. 40, no. 1, pp. 419–430, 2020.

[31] M. Ghandehari, J. Janssen, and N. Kalyaniwalla, “A noncommutative approach to the graphon Fourier transform,” Applied and Computational Harmonic Analysis, vol. 61, pp. 101–131, 2022.

